# Protection against neonatal respiratory viral infection via maternal treatment during pregnancy with the benign immune training agent OM-85

**DOI:** 10.1101/2021.03.02.433517

**Authors:** Jean-Francois Lauzon-Joset, Kyle T Mincham, Naomi M Scott, Yasmine Khandan, Philip A Stumbles, Patrick G Holt, Deborah H Strickland

## Abstract

**Objectives:** Incomplete maturation of immune regulatory functions at birth are antecedent to the heightened risk for severe respiratory infections during infancy. Our forerunner animal model studies demonstrated that maternal treatment with the benign microbial-derived immune modulating agent OM-85 during pregnancy promotes accelerated maturation of immune regulatory networks in the developing fetal bone marrow. Here, we aimed to establish proof-of-concept that this would enhance resilience to severe early life respiratory viral infection during the neonatal period.

**Methods:** Pregnant BALB/c mice were treated orally with OM-85 during gestation and their offspring infected intranasally with a mouse-adapted rhinovirus (vMC_0_) at postnatal day 2. We then assessed clinical course, lung viral titres and lung immune parameters to determine whether offspring from OM-85 treated mothers demonstrate enhanced immune protection against neonatal vMC_0_ infection.

**Results:** Offspring from OM-85 treated mothers display enhanced capacity to clear an otherwise lethal respiratory viral infection during the neonatal period, with a concomitant reduction in the exaggerated nature of the ensuing immune response. These treatment effects were associated with accelerated postnatal myeloid cell seeding of neonatal lungs and enhanced expression of microbial sensing receptors in lung tissues, coupled in particular with enhanced capacity to rapidly expand and maintain networks of lung dendritic cells expressing function-associated markers crucial for maintenance of local immune homeostasis in the face of pathogen challenge.

**Conclusion:** Maternal OM-85 treatment may represent a novel therapeutic strategy to reduce the burden, and potential long-term sequlae, of severe neonatal respiratory viral infection by accelerating development of innate immune competence.

## Introduction

Acute lower respiratory infections (LRI) represent the leading cause of mortality among children younger than 5 years, accounting for more than 15% of all deaths in this age group (1). When compared to the approximate 4.4% of deaths in people of all ages attributable to LRIs each year (2), the urgent requirement for novel treatment strategies to protect specifically against the life-threatening effects of early postnatal respiratory infections becomes immediately apparent. Further reinforcing this need, less severe grades of LRI during infancy are also recognised as a major risk factor for subsequent early onset asthma diagnosis, which contributes to a significant global health and economic burden impacting over 300 million individuals (3–5).

There are major challenges associated with protection of the highly vulnerable neonatal/infant age group from infectious disease mortality. For one, infant responsiveness to immune-enhancing therapeutics, including conventional vaccination, is constrained to variable degrees by developmental-associated deficiencies in innate and adaptive immune functions that mature postnatally. In the present study, we have examined the novel hypothesis that pre-programming key aspects of innate immune function before birth will provide a mechanism to circumvent early developmental immune deficiency, potentially mitigating the risk of death following an early life LRI (6). This approach is based on our forerunner studies involving the microbial-derived immune training agent OM-85, which has been in widespread clinical use for more than 30 years for limiting the risk of severe respiratory infections in young children from preschool age (6–10). Our previous studies using OM-85 treatment of pregnant mice as a preventative strategy have demonstrated that treated mothers developed markedly enhanced resistance to the pregnancy-threatening effects of exposure to bacterial and viral pathogens, which was associated with the modulation of myeloid cell recruitment/egression within inflammed gestational tissues (11). Moreover, OM-85 treatment of pregnant mice was also shown to promote maturation of fetal bone marrow myeloid progenitor populations, resulting in accelerated postnatal establishment of functionally mature dendritic cell (DC) networks in the respiratory mucosal tissues of their offspring. This was accompanied by enhanced capacity of regulatory T-cells (Treg) to regulate the intensity/duration of allergic airways inflammatory responses (12, 13). Given that similar DC/Treg-associated mechanisms are central to protection against infection-induced inflammatory tissue damage in the lower respiratory tract, we posited that maternal OM-85 treatment could also enhance resistance to a lethal viral LRI in the offspring during the highly vulnerable neonatal period, and testing this hypothesis was the focus of the experiments reported below.

## Results

### Accelerated postnatal development of the myeloid compartment within neonatal lungs following maternal OM-85 treatment

Given our previous findings of enhanced myelopoiesis in fetal bone marrow following OM-85 treatment of pregnant mice (12), we first sought to determine the influence of maternal OM-85 treatment on baseline postnatal development of immune cell populations in the lungs of their offspring. Time-mated pregnant mice were orally treated with OM-85 from gestation day (GD) 0.5-17.5, followed by natural term delivery of offspring 2-3 days later (≈GD20.5). Offspring lungs were then harvested and analyzed using multicolour flow cytometry (gating strategies shown in Fig. S1) at postnatal day (PND) 3, 7, 12 and 22. As shown in Fig. 1, the most profound treatment effects were observed during the neonatal/early infancy period (PND 3-7), including accelerated accumulation of plasmacytoid DC (pDC; Fig. 1A), macrophage subsets (Fig. 1B and C) and monocytes (Fig. 1D). We further observed OM-85-induced changes in conventional DC (cDC) 1 and cDC2 populations, with a significant increase in the proportion of both subsets at PND 12 and upregulation of the function-associated molecules MHC class II (I-A/I-E) and CD86 at PND 3-7 (Fig. 1E-J). These OM-85 treatment associated changes in myeloid cell populations were accompanied by enhanced accumulation of myeloid-derived suppressor cells (MDSC; Fig. 1K) which have previously been shown to play a key role in controlling inflammation in neonatal humans and mice (14), and also enhanced accumulation of Treg (Fig. 1L) and cells expressing a phenotype resembling that of type 2 innate lymphoid cells (ILC2; data not shown) at PND 7. Collectively, these findings are consistent with maternal OM-85 treatment transplacentally inducing an enhanced rate of postnatal respiratory myeloid cell maturation in the offspring, with the potential for increased capacity to maintain respiratory immunological homeostasis in the face of proinflammatory challenges during the early postnatal period.

**Fig. 1.**
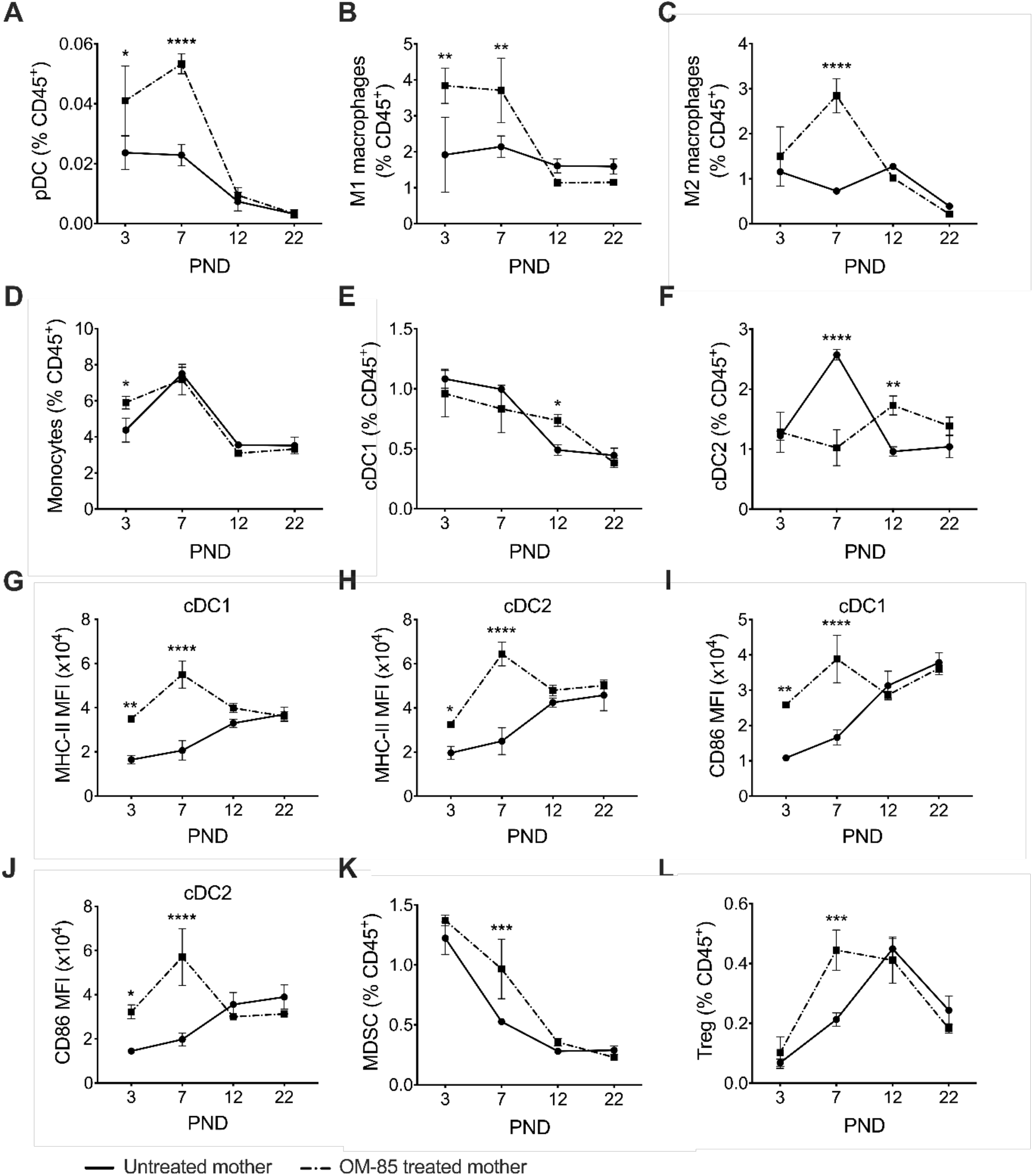
Maternal OM-85 treatment promotes myelopoiesis within neonatal lungs. **(A)** plasmacytoid dendritic cells (pDC), **(B)** M1 macrophages, **(C)** M2 macrophages, **(D)** monocytes, **(E)** conventional DC (cDC) 1 and **(F)** cDC2 as a proportion of CD45^+^ cells within neonatal peripheral lungs. **(G-H)** Mean fluorescence intensity (MFI) of I-A/I-E expression on **(G)** cDC1 and **(H)** cDC2 within neonatal peripheral lungs. **(I-J)** MFI of CD86 expression on **(I)**cDC1 and **(J)**cDC2 within neonatal peripheral lungs. **(K)** Myeloid derived suppressor cells (MDSC) and **(L)** regulatory T-cells (Treg) as a proportion of CD45^+^ cells within neonatal peripheral lungs. Data are displayed as line plot showing mean ± SEM of n = 3-6 offspring from n = 3 independent experiments per group. Statistical significance was determined using two-way ANOVA followed by Uncorrected Fisher’s LSD test; *p < 0.05, **p < 0.01, ***p < 0.001, ****p < 0.0001.

### Maternal OM-85 treatment upregulates TLR4 and TLR7 expression in offspring lungs

To further characterise the underlying mechanism(s)-of-action of maternal OM-85 driven transplacental innate immune enhancement, we measured early postnatal gene expression of a range of immunoregulatory cytokines and cellular sensors within peripheral lungs of the offspring of OM-85-treated mothers. Maternal OM-85 treatment resulted in accelerated postnatal upregulation of both *interleukin (IL)-1α* and *IL-1β* gene expression, which reached statistical significance by PND 22 (Fig. 2A and B), thus mirroring prior observations in independent model systems following OM-85 treatment of murine macrophages (15). Given the dependency of IL-1β production on NOD-like receptor family pyrin domain containing 3 (NLRP3) and caspase-1, we next assessed the impact of maternal OM-85 treatment on gene expression levels of these upstream targets in offspring lungs. As demonstrated in Fig. 2C, maternal OM-85 treatment upregulated *NLRP3* expression at all timepoints measured during the first 22 days of life in their offspring, however downstream concomitant upregulation of *caspase-1* was only evident at PND 22 (Fig. 2D), corresponding to the observed upregulation of *IL-1α/β* expression. Lastly, there is now compelling evidence in both humans (16–18) and experimental murine models (19) demonstrating that the immunomodulatory effects of beneficial microbial exposures, including that of OM-85 (20), are mediated via the enhanced expression of Toll-like receptors (TLRs). We therefore assessed the expression levels of *TLR4* and *TLR7* within neonatal peripheral lungs, with both receptors significantly upregulated across all early postnatal timepoints within offspring from OM-85 treated mothers (Fig. 2E and F), consistent with the centrality of this innate mechanism in immunomodulator-mediated transplacental immune training.

**Fig. 2.**
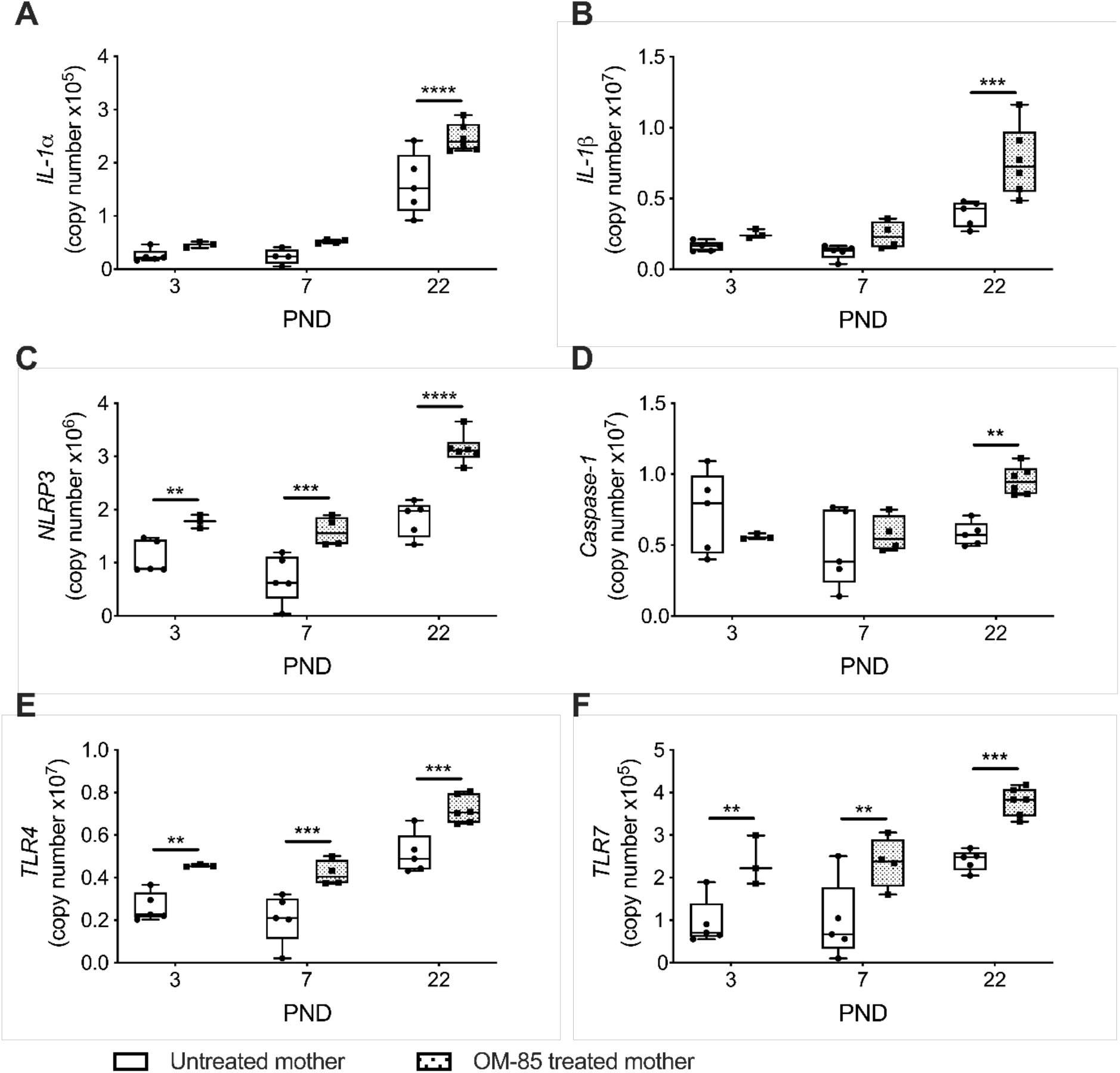
Maternal OM-85 treatment upregulates the IL-1β pathway and TLR expression in neonatal lungs. Absolute copy number of gene expression profiles for **(A)** *IL-1α*, **(B)** *IL-1β*, **(C)** *NLRP3*, **(D)** *Caspase-1,* **(E)** *TLR4* and **(F)** *TLR7* within the lungs of neonates from OM-85 treated and untreated mothers. Data are from individual mice and displayed as box and whisker plot showing minimum to maximum values of n = 3-6 offspring from n = 3 independent experiments per gene. Statistical significance was determined using two-way ANOVA followed by Uncorrected Fisher’s LSD test; **p < 0.01, ***p<0.001, ****p < 0.0001.

### Maternal OM-85 treatment protects neonates against vMC_0_-induced death

Having demonstrated that maternal OM-85 treatment enhanced postnatal establishment of respiratory immunocompetence, as defined by accelerated myelopoiesis and innate cytokine signalling pathways, we next sought to determine if this would translate into enhanced protection against a lethal neonatal LRI. To address this, we developed a murine neonatal infection model of human rhinovirus (HRV), using live attenuated Mengovirus (vMC_0_), a mouse-adapted mimic of HRV (21). Initially, neonatal offspring of untreated mothers were infected with titrated doses of vMC_0_ at PND 2 (henceforth refered to as day-post-infection [DPI] 0), and survival and weight loss assessed up to DPI 20 (Fig 3A and B). Given the age-dependent response observed in murine respiratory viral infection models (22–24), our initial experiments aimed to determine the appropriate vMC_0_ dose required to achieve 20%-25% death (LD_20-25_) in PND 2 neonates. The typical experimental adult mouse LD_20-25_ dose of 10^6^ plaque-forming-units (PFU) (21) resulted in ≈90% death of infected PND 2 neonates by DPI 12 (Fig. 3A). With further titration, an LD_20-25_ dose in neonates was achieved following infection with 10^4^ PFU vMC_0_, and this dose was used for the remainder of the study (Fig. 3A). Neonates infected with 10^4^ PFU vMC_0_ began showing signs of weight loss from DPI 8 when measured as a ratio to uninfected litter mates, with maximal weight loss by DPI 18 and partial recovery by DPI 20 for surviving mice (Fig. 3B). We next sought to determine if the severity of vMC_0_-induced clinical disease in neonates could be modulated following OM-85 treatment of the mothers during gestation, as described above (Fig. 3C). As illustrated in Fig. 3D, maternal OM-85 treatment resulted in 100% protection against vMC_0_-induced neonatal death when compared to the offspring of non-treated mothers, while weight loss in vMC_0_ infected neonates was significantly less severe in those from OM-85 treated mothers compared to surviving infected neonates from untreated mothers (Fig. 3E). Furthermore, maternal OM-85 treatment during pregnancy lead to significantly reduced peak viral titres in both the lungs and brain of vMC_0_ infected neonates at the peak of infection compared to the offspring of untreated mothers (Fig. 3F and G), consistent with our previous findings demonstrating enhanced viral clearance in OM-85 treated versus untreated mothers (11) and independent studies in adult mice (25).

**Fig. 3.**
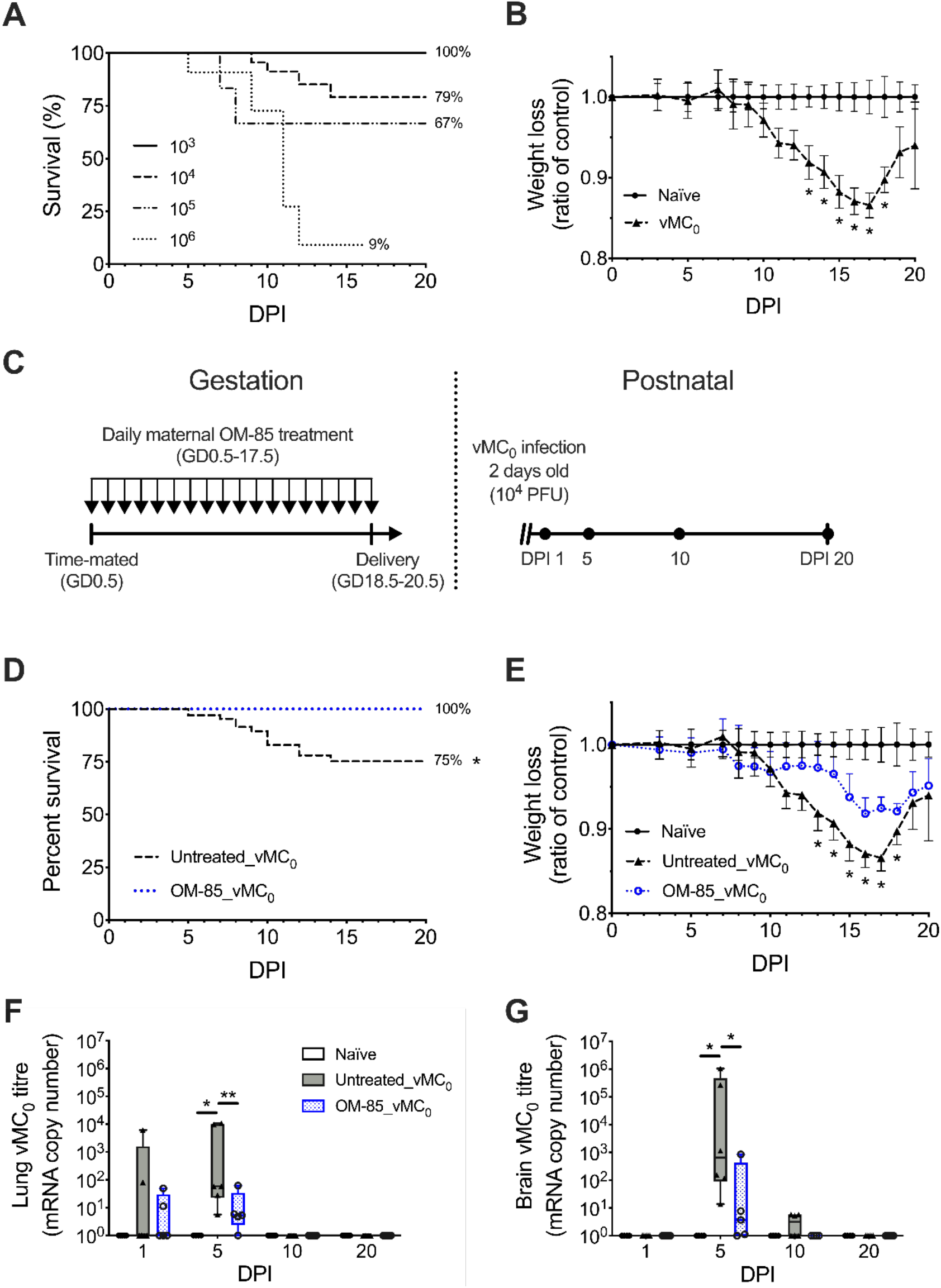
Maternal OM-85 treatment protects neonates against vMC_0_ respiratory viral infection-induced death. **(A)** Titration of neonatal vMC_0_ plaque-forming-units (PFU) required to achieve a lethal dose in 20%-25% (LD_20-25_) of infected neonates (10^3^ *n* = 12, 10^4^ *n* = 38, 10^5^ *n* = 6, 10^6^ *n* = 11). Data are displayed as survival curves showing n = 12 independent experiments. **(B)** Weight loss of neonates infected with 10^4^ PFU (Naïve *n* = 28, vMC_0_ *n* = 33;). Data are displayed as line plot showing mean ± SEM of n = 8 independent experiments. Statistical significance was determined using two-way ANOVA with Tukey multiple comparisons test; *p < 0.05. **(C)** Maternal OM-85 treatment regimen from gestation day (GD) 0.5-17.5 (left) followed by neonatal vMC_0_ infection at 2 days of age and disease kinetics monitored until day post infection (DPI) 20 (right). Neonates were autopsied and samples collected at DPI 1, 5, 10 and 20. **(D)** Survival curves of vMC_0_ infected neonates from OM-85 treated (OM-85_vMC_0_; *n* = 19) and untreated (Untreated_vMC_0_; *n* = 33) mothers. Data are displayed from n = 8 independent experiments. Statistical significance was determined using Log-rank (Mantel-Cox) test and displayed as *p < 0.05. **(E)** Weight loss of vMC_0_ infected neonates from OM-85 treated and untreated mothers (Naïve *n* = 28, Untreated_vMC_0_ *n* = 33, OM-85_vMC_0_ *n* = 19). Data are displayed as line plot showing mean ± SEM of n = 12 independent experiments. Statistical significance was determined using two-way ANOVA with Tukey multiple comparisons test; *p < 0.05 vs. control. **(F-G)** vMC_0_ viral titres in the **(F)** lungs and **(G)** brain of neonates from OM-85 treated (OM-85_vMC_0_) and untreated (Untreated_vMC_0_) mothers. vMC_0_ viral load of the control neonates was below the detection limit of the assay. Data are displayed as box and whisker plot showing minimum to maximum values of n = 4-6 offspring from n = 3 independent experiments. Statistical significance was determined using two-way ANOVA followed by Uncorrected Fisher’s LSD test; *p < 0.05.

### Attenuation of vMC_0_-induced innate proinflammatory cellular responses in neonatal lung tissue following maternal OM-85 treatment

In addition to the clinical assessments above, multicolour flow cytometry was performed on single cell lung preparations to determine the impact of maternal OM-85 treatment on the cellular immune response to neonatal vMC_0_ infection. Immunophenotypic characterization of neonatal lungs revealed that the severity of the classical acute proinflammatory innate viral response was significantly attenuated in those neonates from OM-85 treated mothers compared to equivalent neonates from untreated mothers. This included a reduced influx of neutrophils at DPI 1 (Fig. 4A), natural killer (NK) cells at DPI 1 and DPI 5 (Fig. 4B), and classical monocytes over the entire disease time course evaluated (Fig. 4C). The early influx of pDC into the lungs of vMC_0_ infected neonates at DPI 1 (Fig. 4D) was also dramatically less intense in offspring from OM-85 treated compared to untreated mothers, however this pDC population exhibited a substantial increase in expression of MHC class II at the peak of viral load (Fig. 4E) and CD40 expression during the ensuing resolution phase (Fig. 4F). Immunophenotypic characterization additionally identified multiple lung macrophage subsets in neonates, however maternal OM-85 treatment had very little influence on these cell populations in vMC_0_ infected neonates compared to offspring of untreated mothers (Fig. S2A-C). Together, these data indicate that maternal OM-85 treatment induced a shift in the lung cellular immune response of the offspring following neonatal vMC_0_ infection towards a state of improved homeostatic control and dampened inflammation within the lungs, while at the same time reducing viral load.

**Fig. 4.**
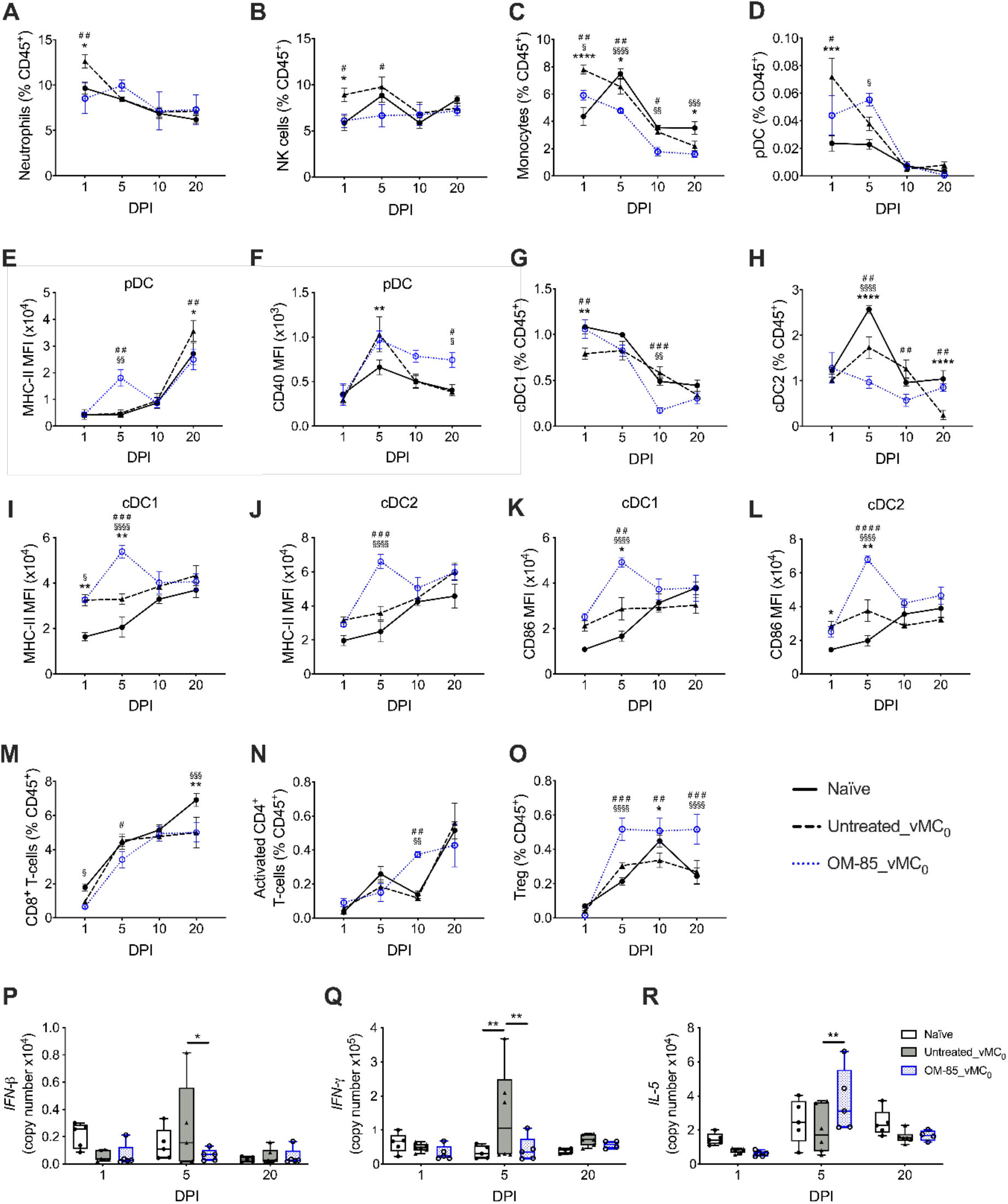
Maternal OM-85 treatment dampens the innate proinflammatory cellular response whilst enhancing regulatory responses within neonatal lungs following vMC_0_ infection. **(A)** Neutrophils, **(B)** NK cells, **(C)** monocytes and **(D)** pDC as a proportion of CD45^+^ cells within neonatal peripheral lungs. **(E-F)** MFI of **(E)** I-A/I-E and **(F)** CD40 expression on pDC. **(G)** cDC1 and **(H)** cDC2 as a proportion of CD45^+^ cells. **(I-J)** MFI of I-A/I-E expression on **(I)** cDC1 and **(J)** cDC2. **(K-L)**MFI of CD86 expression on **(K)** cDC1 and **(L)** cDC2. **(M)** CD8^+^ T-cells, **(N)** activated CD4^+^ T-cells and **(O)** Treg as a proportion of CD45^+^ cells. Data are displayed as line plot showing mean ± SEM of n = 4-6 offspring from n = 4 independent experiments per group. Statistical significance was determined using two-way ANOVA followed by Uncorrected Fisher’s LSD test; *Naïve vs. Untreated_vMC_0_, ^§^Naïve vs. OM-85_vMC_0_, ^#^Untreated_vCM0 vs. OM-85_vMC_0_. **(P-R)** Absolute copy number of gene expression profiles for **(P)** *IFN-β*, **(Q)** *IFN-γ* and **(R)** *IL-*5 within neonatal peripheral lungs. Data are displayed as box and whisker plot showing minimum to maximum values of n = 4-6 offspring from n = 3 independent experiments per gene. Statistical significance was determined using two-way ANOVA followed by Uncorrected Fisher’s LSD test; *p < 0.05, **p < 0.01.

Within the lungs, cDC subsets represent a highly dynamic population that turn over rapidly, providing continuous signalling of incoming airborne antigens impacting respiratory surfaces to the T-cell system in lymph nodes and respiratory tissues (26, 27). In this regard, cDC1 are primarily responsible for the cross-presentation of viral antigen to CD8^+^ T-cells (26, 28), while their cDC2 counterparts are primariliy involved in the generation of CD4^+^ T-cell responses within regional airways draining lymph nodes. Following viral challenge of neonates from untreated mothers, we observed the specific modulation of both cDC subsets in neonatal lung tissue, including a significant reduction in cDC1 at DPI 1 (Fig. 4G), while cDC2 proportions were reduced at the peak of infection at DPI 5 (Fig. 4H), likely representing the complex interplay between cellular efflux/influx within the bone marrow-lung-lymph node axis (29). Of particular interest, vMC_0_ infected neonates from untreated mothers exhibited a deficit in cDC2 numbers which persisted until infection resolution at DPI 20 (Fig. 4H), consistent with previous findings that early life influenza infection drives persistent dysregulation of local myeloid populations (30). However, maternal OM-85 treatment was able to preserve lung cDC2 numbers at similar levels to those observed in age matched non-infected controls (Fig 4H). Moreover, both cDC1 and cDC2 subpopulations displayed enhanced functional upregulation in the lungs of neonates from OM-85 treated mothers in response to vMC_0_ challenge as evidenced by a significant increase in the function-associated molecules MHC class II (Fig. 4I and J) and CD86 (Fig. 4K and L) at peak viral load at DPI 5, although with no observable changes in CD40 expression (Fig. S2D and E). Collectively these observations are consistent with previous findings (13) indicating that maternal OM-85 treatment transplacentally regulates the fetal/neonatal bone marrow-lung axis towards development of lung myeloid cell regulatory networks more resilient to excessive inflammation in the face of early life respiratory inflammation, through enhanced developmental trajectories of lung innate immune cell subsets that are crucial for maintenance of local immune homeostasis.

We additionally assessed T-cell responses in the lungs of vMC_0_ infected neonates from OM-85 treated and untreated mothers. At the peak of clinical disease (DPI 5), maternal OM-85 treatment induced a small but significant reduction in the frequency of CD8^+^ T-cells in the lungs of vMC_0_ infected offspring compard to equivalent offspring of untreated mothers (Fig. 4M). Notably, while there was no significant influence of maternal OM-85 treatment on the overall CD4^+^ T-cell pool in vMC_0_ infected neonates (Fig. S2F), we identified an enhanced population of activated CD4^+^ T-cells at DPI 10 in infected neonates from OM-85 treated mothers (Fig. 4N). These same neonates additionally displayed a significantly amplified pool of mucosal homing Treg within the peripheral lung from the peak of disease until resolution at DPI 20 (Fig. 4O), while cells resembling a local ILC2 population were significantly upregulated from DPI 1-5 in vMC_0_ infected neonates from OM-85 treated (data not shown). These findings are consistent with our previous studies and those of others identifying Tregs as a crucial target for both the direct (20, 31) and transplacental (13) protective effects of OM-85 treatment in inflammation-induced respiratory diseases during early life.

Finally, to gain further insight into the mechanisms promoting transplacental OM-85-mediated protection against vMC_0_ infection severity in neonatal offspring, we measured the gene expression levels of key cytokines in neonatal lungs, including Type I and Type II IFNs typically associated with anti-viral responses. In line with reduced viral titres and clinical disease severity at the peak of infection in neonates from OM-85 treated mothers, we observed significantly dampened IFN responses in vMC_0_ infected neonates when compared to those from untreated mothers, as evidenced by a reduction in both *IFN-β* (Fig. 4P) and *IFN-γ* (Fig. 4Q) gene expression. Moreover, neonates of mothers exposed to gestational OM-85 treatment additionally displayed a heightened type 2-associated *IL-5* gene response when compared to those from untreated mothers (Fig. 4R).

## Discussion

Acute LRIs are the leading cause of death among children below the age of 5 years, with risk being highest in the neonatal/infancy period that precedes completion of the postnatal development process leading to establishment of immune competence. In the animal model studies presented here, we tested the hypothesis that mitigation of this risk may be achievable by stimulation of precocious postnatal maturation of immune competence via maternal treatment with a bacterial-derived immune training agent (OM-85) during pregnancy. As a proof-of-concept, our data demonstrate that maternal OM-85 treatment significantly protects the offspring against an LD_20-25_ dose of the HRV mimic vMC_0_ given during the early neonatal period. Consistent with previous infection-related studies examining the effects of oral OM-85 treatment in adult mice (11, 25), protection was associated with significant attenuation of viral load within the lungs, and for the first time we demonstrate parallel reduction of viral load within the brain of vMC_0_ infected neonates. It is pertinent to note in this regard that viral-related acute encephalopathy is one of the leading complications associated with viral-induced mortality in children (32, 33). At the cellular level, we demonstrate significant attenuation of the acute innate response within the peripheral lungs of vMC_0_ infected neonates from OM-85 treated mothers. Moreover, consistent with our previous reports in an experimental model of allergic airways inflammation (13), maternal OM-85 treatment enhanced the early postnatal expression of key function-associated molecules (MHC class II and CD86) on airway cDC subsets within vMC_0_ infected neonates. This cellular response was accompanied by concomitant attenuation of the intensity of *IFN* gene expression profiles in vMC_0_ infected neonatal lungs at the peak of disease, whilst boosting type 2-associated *IL-5* gene expression.

Within both humans and experimental murine models, the classical pathophysiology of an innate anti-viral response within the airways is characterized by the rapid influx of inflammatory neutrophils into the lungs, intially from the intra-pulmonary endothelial marginating pool and subsequently from the bone marrow via the circulating pool (21, 34, 35), with concurrent secretion of uniquely high levels of Type I IFN-β and Type II IFN-γ by primarily pDC and NK cells respectively (36, 37). However, human studies indicate that persistent activation of these innate pathways and aberrant/dysregulated production of inflammatory cytokines can lead to collateral tissue damage within the lungs and subsequent chronic lung pathology or even death (38–40). This disadvantageous inflammatory response is most prominent in infants with acute viral bronchiolitis, whereby those presenting with severe symptoms, including febrile episodes, display hyperactivation of IFN-associated gene networks within both whole blood/peripheral blood mononuclear cells and the nasal mucosa (41, 42). Our current findings demonstrating enhanced regulation of the intensity of *IFN* responses to vMC_0_ infection in neonatal mouse lungs mirrors those in our earlier study on the mechanism-of-action of OM-85 in protection of maternal gestational challenge against the toxic effects of bacterial challenge, in which the main target of OM-85 treatment was identified as proinflammatory IFN-γ gene networks (11). It is additionally pertinent to note that suppression of IFN-γ during murine respiratory viral infection has been reported to enhance ILC2 function in the lungs (43), which in conjunction with the upregulated subset of cells displaying an ILC2-like phenotype at DPI 5, provides a possible explanation as to the cellular source of enhanced *IL-5* gene expression within the lungs of vMC_0_ infected neonates from OM-85 treated mothers, given their capacity to produce greater amounts of IL-5 than CD4^+^ T-cells on a per-cell basis following antigenic stimulation (44). Moreover, these observations are in line with our forerunner studies identifying enhanced T-cell activation-associated signatures in fetal bone marrow following maternal OM-85 treatment during pregnancy (12). In regards to T-cell signatures, the enhanced pool of Treg within the lungs of vMC_0_ infected offspring from OM-85 treated mothers is consistent with previous findings by ourselves (13, 31) and others (20) whereby oral OM-85 treatment boosts the airways Treg pool in response to local inflammatory challenge, and parallels independent studies demonstrating a role for Treg in the clearance of respiratory virus in neonatal mice (45).

Insight into the nature of OM-85-induced transplacental mechanisms promoting offspring protection to respiratory viral infection was provided in the offspring of OM-85 treated mothers in the absence of neonatal vMC_0_ infection, whose lungs displayed an enriched baseline pool of cDC subsets as evidenced by their increased numbers and maturation/activation states. In this regard it is pertinent to note that during this early postnatal period, immunoregulatory DC networks in the airway mucosa are typically developmentally compromised in both humans (46, 47) and experimental animals (48–50) in regards to both population density and functional maturity. These findings mirror our previous discovery of accelerated postnatal establishment of functionally mature DC networks in respiratory tissues of offspring from OM-85 treated mothers (13). Furthermore, we have previously identified the capacity of OM-85 to act transplacentally to expand the fetal bone marrow myeloid progenitor compartment by late gestation (12), which ultimately gives rise to the broad repertoire of DCs responsible for seeding the lungs during postnatal immunological development. Concommitent with the enhanced innate myeloid response, maternal OM-85 treatment additionally enriched the baseline pool of lung Tregs in neonates at 7 days of age, further supporting a role for transplacental OM-85 treatment in the accelerated establishment of a basal regulatory state in neonatal lungs.

Studies performed by our research team (12) have previously identified analogous hallmark immune training signatures shared between OM-85 treatment and the classical training agents β-glucan and BCG (51–53). In the data presented here, we further expand on these shared features by identifying an additional parallel mechanism in the form of upregulated IL-1β responsiveness (51). IL-1β has primarily been investigated for its anti-viral properties, a process dependent on stimulation of the NLRP3 inflammasome and resultant activation of caspase-1 (54–56). Moreover, pre-treatment with recombinant IL-1β has proven to protect adult mice against lethal *Pseudomonas aeruginosa* infection, thus demonstrating anti-bacterial capabilities (57). However, studies have now identified the pleotropic role of IL-1β in promoting proliferation and differentiation of myeloid progenitors (51, 58), which have recently been isolated from murine lungs (59, 60), thereby fostering a tissue microenvironment tailored towards myelopoiesis in the absence of inflammatory perturbation. Likewise, activation of the NLRP3 inflammasome and downstream signalling via IL-1 has recently been recognised as a mechanism of innate immune training in response to dietary influences, complementing previously identified mechanisms via promoting the expansion of myeloid progenitor subsets (61). Within the data presented here, the dual NLRP3/caspase-1 signal required for IL-1β expression is only observed within offspring lungs at PND 22 following maternal OM-85 treatment, paralleling previous studies revealing that OM-85 acts as a priming signal for the NLRP3 inflammasome complex (62), and suggests that optimisation of the peripheral lung myeloid network is maintained throughout later life. In this regard, we have now established that OM-85-mediated upregulation of offspring myelopoiesis occurs from *in utero* development into adolescence (12, 13), thereby demonstrating that multiple mechanisms, operating over distinct time scales, may be responsible for this enhanced functional immunocompetence.

In this regard, a crucial mechanistic finding within the study presented here is the enhanced TLR expression in neonatal peripheral lung following maternal OM-85 treatment. Differentiation of myeloid cells is dependent on signalling via TLRs (63–65), with impaired TLR signalling during early life development in-part responsible for the functional immaturity of the myeloid network within neonates (66). Moreover, levels of TLR expression are recognised as key determinants of responsiveness to environmental microbial stimuli exemplified by farm dust which are known to promote development of resistance to early asthma onset in both humans (16, 17, 67) and murine models (19). Specific to respiratory viral infection, pre-exposure of human primary bronchial epithelial cells to farm dust extract significantly enhances TLR2 expression, and limits HRV viral load in a TLR2/barrier function-dependent manner (18), while neonatal murine TLR4 stimulation prior to severe RSV infection promotes the generation of an “adult-like” anti-viral response, associated with an increase in cDC activation within the airways (68), mirroring features of the data presented here. Taken together, we therefore postulate that optimisation of innate microbial sensing machinery via enhanced TLR4 and TLR7 expression within neonatal peripheral lungs is one of the primary maternal OM-85-mediated mechanisms promoting upregulation of myeloid subsets and accelerated functional immunocompetence of the neonatal respiratory myeloid network. Furthermore, we have previously identified the maternal OM-85-mediated modulation of microRNAs associated with TLR4 expression within fetal bone marrow (12), signifying that the enhanced offspring peripheral lung response may be a direct result of immune training events initiated *in utero*.

We acknowledge several inherent limitations in this study which need to be addressed in follow-up investigations. Firstly, we have no information on the cellular/transcriptomic response occurring between DPI 1 and 5, and such data may provide valuable insight into which neonates would ultimately succumb to viral infection and which would be protected, given the transient nature of the innate viral response following acute vMC_0_ infection (21, 69). Secondly, we are unable to make precise distinctions between the roles of immune/inflammatory cell populations which have extravasated into peripheral lung tissues and corresponding marginal pools adherent to vessel walls, given the inability to efficiently perfuse fragile neonatal lungs prior to tissue harvest. In this regard it is known that the marginated pool of lung immune/inflammatory cells, particularly neutrophils (35, 70), is primed to rapidly extravasate from the lung vasculature and thus represents a crucial component of the acute innate response to inflammatory stimuli, and the lung-associated peripheral T-cell compartment is similarly partitioned (71, 72); this particular limitation is thus common to almost all studies in this area. Lastly, we have not identified the cellular source within lung tissue of enhanced TLR expression in the offspring of OM-85 treated mothers. In this regard, mesenchymal and epithelial cells within the lungs are known to express a broad repertoire of TLRs crucial in dictating function during immune responses (73, 74). Moreover, the process of trained immunity has recently been extended beyond hematopoietic immune cells (18, 75). The potential solution to these issues in the future may be to employ a combination of cell sorting/single-cell RNA sequencing to enable high-precision mapping of the innate response on a per-cell basis in different lung tissue compartments. Notwithstanding these limitations, the strength of the protective effects of OM-85 in this model against respiratory pathogen challenge in the uniquely susceptible neonatal/infant age group provides strong justification for progression towards translational studies in human pregnancy.

## Materials and Methods

### Animals

Specific pathogen-free BALB/c mice were purchased from the Animal Resource Centre (Murdoch, Western Australia). All mice were house under specific pathogen-free conditions with standard food and water *ad libitum* at the Telethon Kids Institute Bioresources Centre.

### Time-mated pregnancies

Female BALB/c mice 8-12 weeks of age were time-mated with male BALB/c studs 8-26 weeks of age. Male studs were individually housed with 1-2 females overnight. The detection of a vaginal plug the following morning was designated gestation day (GD) 0.5.

### Maternal OM-85 treatment regimen

OM-85 (OM Pharma) is an endotoxin-low lyophilised standardised extract containing multiple TLR ligands derived from 8 major common respiratory bacterial pathogens (*Haemophilus influenzae, Streptococcus pneumonia, Streptococcus pyogenes, Streptococcus viridians, Klebsiella pneumoniae, Klebsiella ozaenae, Staphylococcus aureus, Neisseria catarrhalis*) (76). Based on previously optimised dosing concentrations (11, 13), time-mated pregnant BALB/c mice received daily oral feeding of lyophilised OM-85 reconstituted in phosphate-buffered saline (PBS) at a concentration of 400mg/kg body weight from GD0.5-17.5. Control pregnant mice were left untreated for the duration of the study. All maternal treatment was performed with a single batch of OM-85 (batch# 1812162).

### Neonatal Mengovirus (vMC_0_) infection

Attenuated Mengovirus (vMC_0_) was prepared as previously described (21). Two-day-old BALB/c neonatal mice from OM-85 treated and untreated mothers were intranasally inoculated with 10μl of either 10^3^, 10^4^, 10^5^ or 10^6^ plaque-forming-units (PFU) of live attenuated vMC_0_. Following vMC_0_ dose titrations, 10^4^ PFU was used for all experimental inoculations.

### Tissue collection

Neonates were autopsied at day-post-infection (DPI) 1, 5, 10 and 20, which equates to postnatal day (PND) 3, 7, 12 and 22. Pre-perfusion of lung tissues from neonatal/infant animals was not feasible due to tissue fragility and so we standardized throughout to collection of samples from non-perfused lungs, which hence included marginating immune/inflammatory cell populations present on endothelial surfaces of the lung vascular bed (35, 70–72, 77–79), plus a small contribution from occluded blood. Peripheral lung and brain tissue were collected and stored in either cold PBS + 0.1% bovine serum albumin (BSA) for flow cytometric analysis, or RNAlater^®^ stabilization solution (Sigma-Aldrich) for analysis of gene expression profiles and viral titres. Samples collected into RNAlater^®^ were stored overnight at 4°C, then transferred to 1.5ml Eppendorf tubes (Eppendorf) and frozen at −80°C for future analysis.

### Single cell suspension preparation

Neonatal lungs were prepared by enzymatic digestion as previously described (13). Briefly, lungs were minced with a scalpel and resuspended in 10ml GKN + 10% fetal calf serum (FCS) with collagenase IV (Worthington Biochemical Corp.) and DNase (Sigma-Aldrich) at 37°C under gentle agitation for 60 minutes. Digested cells were filtered through sterile nylon, centrifuged and resuspended in cold PBS for total cell counts.

### Flow cytometry

Neonatal lung single cell suspensions were used for all immunostaining. Panels of monoclonal antibodies (purchased from BD Bioscience unless otherwise stated) were developed to enable phenotypic characterization of leukocytes of myeloid: CD45-PerCP (clone 30-F11), CD11b-v500 (clone M1/70), CD11c-AF700 (clone HL3), CD19-BV786 (clone 1D3), CD103-PE (clone M290), CD301-PE-Cy7 (clone MGL1/MGL2. BioLegend), F4/80-FITC (clone BM8; BioLegend), Ly6G/C-APC-Cy7 (clone RB6-8C5), I-A/I-E-AF647 (clone M5/114.14.2), B220/CD45R-PE-CF594 (clone RA3-6B2) and lymphoid: CD45-PerCP (clone 30-F11), NKp46-PE-Cy7 (clone 29A1.4; BioLegend), CD19-BV786 (clone 1D3), CD3-FITC (clone 17A2), CD4-v500 (clone RM4-5), CD8α-BV650 (clone 53-6.7), CD25-APC-Cy7 (clone PC61), Foxp3-PE (clone FJK-16s) lineages. Intracellular staining for Foxp3 was performed using an intracellular Foxp3/Transcription factor staining buffer kit (eBioscience). All samples were kept as individuals. Immune cell phenotypic characterization was performed using FlowJo software (version 10.6.1, BD Bioscience). Fluorescent minus one staining controls were used for all panels where necessary. Flow cytometry data quality was based on primary time gates to ensure appropriate laser delay (pre-determined by automated CS&T) during sample acquisition.

### Viral titre

vMC_0_ viral titre was measured in lung and brain homogenates by real-time quantitative polymerase chain reaction (RT-qPCR). Lungs and brain were homogenised in PBS (10% w/v) using a rotor-star homogeniser (Qiagen) and total RNA extracted using TRIzol (Invitrogen) and RNeasy MinElute Cleanup Kit (Qiagen). Complementary DNA (cDNA) was prepared via QuantiTect Reverse Transcription Kit (Qiagen) and vMC_0_ viral copy number determined using the QuantiFast SYBR Green PCR Kit (Qiagen) via the primer sequence; *forward primer*: 5’-GCC GAAAGC CAC GTG TGT AA and *reverse primer*: 5’-AGA TCC CAG CCA GTG GGG TA (69). Viral copy numbers were calculated using a standard curve of known amounts of amplified cDNA.

### Cytokine and cellular sensor analysis

Cytokines and cellular sensors were measured in neonatal lung homogenates (cDNA prepared as per viral titres) by RT-qPCR using QuantiFast SYBR Green PCR Kit (Qiagen) and QuantiTect Primer Assays (Qiagen; as per manufacturer’s instructions) for the detection of *IL-1α*, *IL-1β*, *IL-5*, *IFN-β*, *IFN-γ*, *NLRP3*, *Caspase-1*, *TLR4* and *TLR7*.

### Statistical analysis

Statistical analysis and graphing was performed using GraphPad Prism (version 8.3.0, GraphPad Software). Statistical significance of p < 0.05 was considered significant. Two-way analyses of variance (ANOVA) followed by Uncorrected Fisher’s LSD test or Tukey’s multiple comparisons test and Log-rank (Mantel-Cox) test were used for analyses as outlined in corresponding figure legends.

### Study approval

All animal experiments were formally approved by the Telethon Kids Institute Animal Ethics Committee, which operates under guidelines developed by the National Health and Medical Research Council of Australia for the care and use of animals in scientific research.

## Acknowledgements

The authors wish to acknowledge the animal technicians at the Telethon Kids Institute Bioresources Centre.

## Funding

This study was funded by the Asthma Foundation of Western Australia and Telethon Kids Institute. JFLJ was supported by a fellowship from the Fond de Recherche du Québec en Santé.

## Author Contributions

JFLJ and DHS designed and supervised the study. KTM and YK performed the experiments. JFLJ, KTM, NMS, PAS, PGH and DHS interpreted the data. JFLJ, KTM, PAS, PGH and DHS wrote and revised the manuscript.

## Competing Interests

All authors declare that no competing interests exist.

## Notes

### Competing Interest Statement

The authors have declared no competing interest.

