## Supplementary Figures for "Protection against neonatal respiratory viral infection via maternal treatment during pregnancy with the benign immune training agent OM-85"

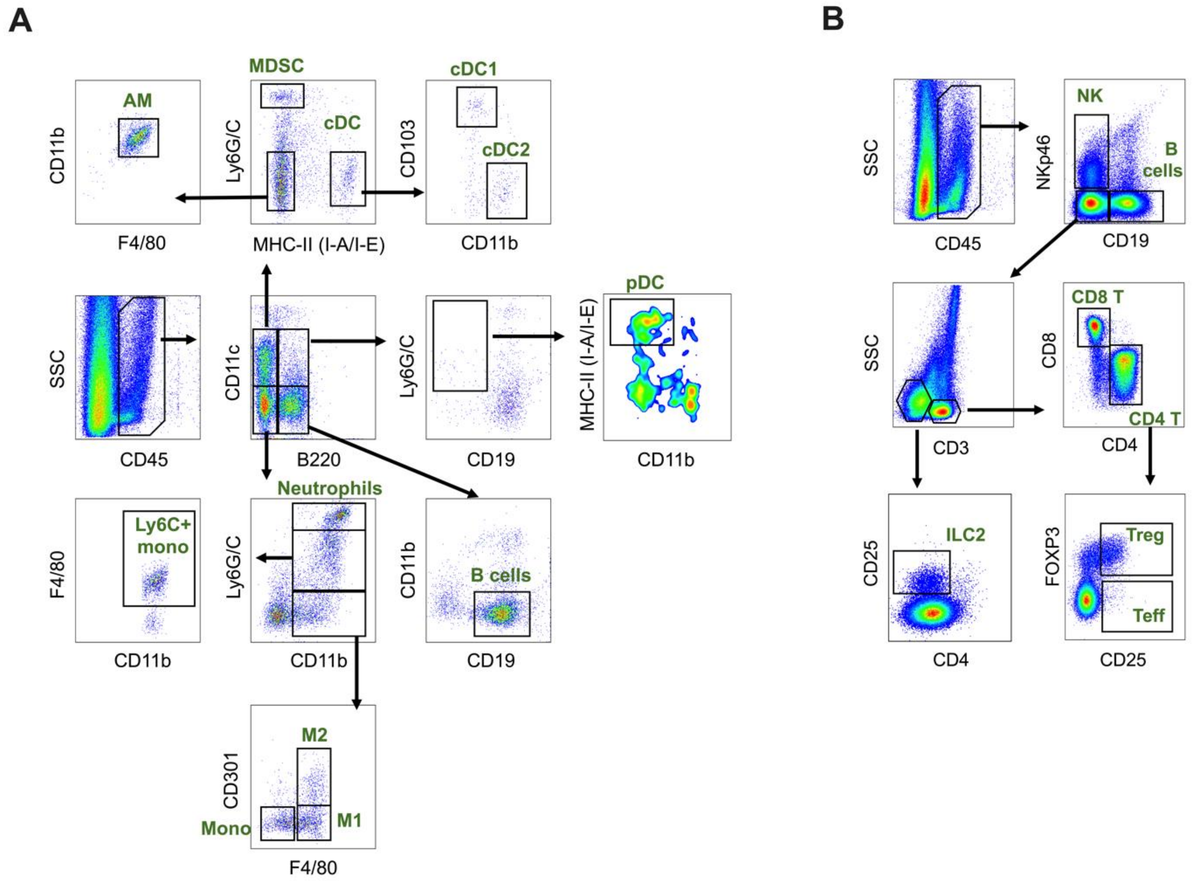


**Fig. S1. Immunophenotypic gating strategies.** Representative FlowJo gating strategies used to identify **(A)** myeloid cells and **(B)** lymphocytes in lung samples via multicolour flow cytometry.

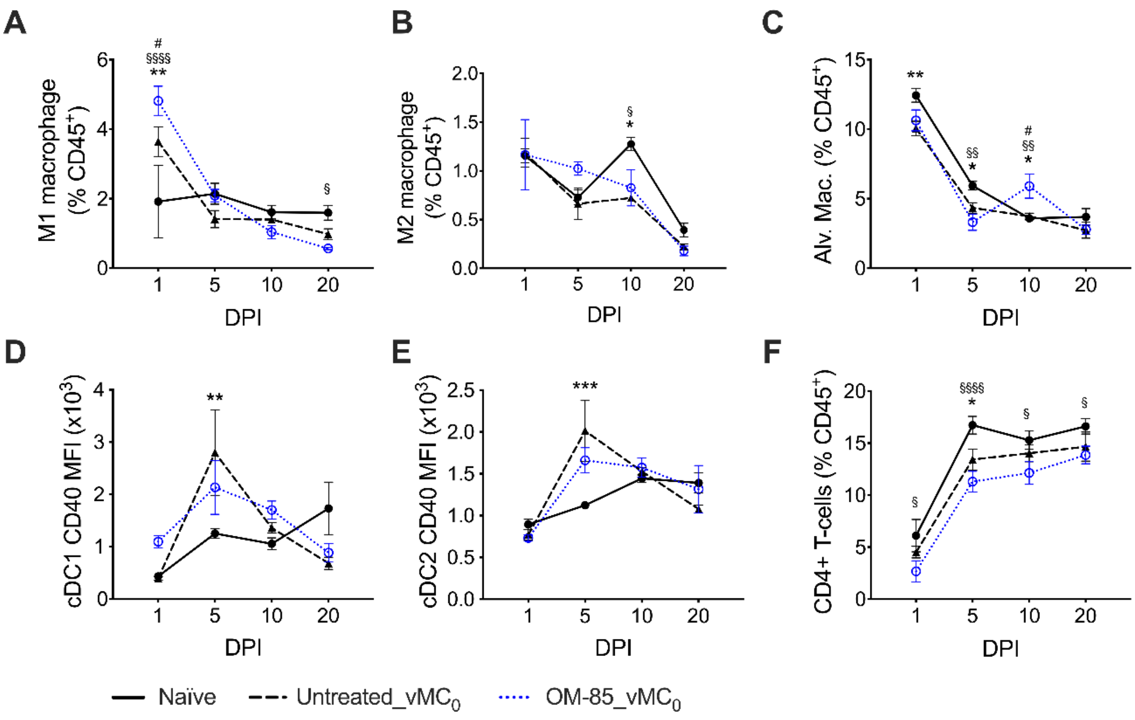


**Fig. S2. Cellular response within $vMC_0$ infected neonates from OM-85 treated and untreated mothers. (A)** M1 macrophages, **(B)** M2 macrophages and **(C)** alveolar macrophages as a proportion of $CD45^+$ cells within neonatal peripheral lungs. MFI of CD40 expression on **(D)** cDC1 and **(E)** cDC2 within neonatal peripheral lungs. **(F)** $CD4^+$ T-cells as a proportion of $CD45^+$ cells within neonatal peripheral lungs. Data are presented as line plots showing mean ± SEM of n = 4-6 offspring from n = 4 independent experiments. Statistical significance was determined using two-way ANOVA followed by Uncorrected Fisher's LSD test; *Naïve vs. Untreated_$vMC_0$, §Naïve vs. OM-85_$vMC_0$, #Untreated_$vCM_0$ vs. OM-85_$vMC_0$.
